# Efficacy of recombinant NP-M1 and NP-M1-CRT DNA vaccines against Influenza A viruses in mice C57/BL6

**DOI:** 10.1101/2021.06.24.449692

**Authors:** Hamidreza Attaran, Wen He, Wei Wang

## Abstract

Effective vaccination against the influenza virus remains a challenge because of antigenic shift and drift in influenza viruses. Conservation is an important feature of the Nucleoprotein (NP) and Matrix protein 1(M1) qualifying them as potential candidates for developing a universal vaccine against the influenza A virus. Carliticulin (CRT), a member of heat shock protein (HSP) family, are conserved and widely distributed in many microorganisms and mammalian cells. In this study, a plasmid vector encoding the NP-M1-CRT sequence was constructed and compared with the NP-M1 sequence with respect to immunogenicity and protective efficacy in a murine model. The potency of the created construct for provoking humoral, cellular immune responses, and its protective immunity against the lethal influenza virus infection were then compared with commercial split vaccine and then evaluated in a murine model system. NP-M1-CRT as a DNA vaccine combined with in vivo electroporation could significantly improve the immunogenicity of constructed vectors. Serological evaluations demonstrated the potency of our approach to provoke strong anti-NP specific antibody responses. Furthermore, our strategy of immunization in prime-boost groups were able to provide protection against lethal viral challenge using H1N1 subtype. The ease of production of these types of vectors and the fact that they would not require annual updating and manufacturing may provide an alternative cost-effective approach to limit the spread of potential pandemic influenza viruses.

## 1. Introduction

The influenza viral infection is one of the most common and widespread viral diseases in mammalian and avian species. Often, a virulent strain can be lethal and may take a catastrophic toll if preventive measures are not taken. Annual vaccination is an important preventative measure that reduces morbidity and mortality rates to the virus [1]. Today, licensed influenza vaccines primarily induce strain-specific antibodies to hemagglutinin (HA) and neuraminidase (NA) [2,3]. There are several major setbacks with the current model utilized by licensed influenza A vaccine producers: Namely, the requirement to produce new vaccines every season, the uncertainty in choosing the accurate influenza A virus strain to create a vaccine for that season, long manufacturing cycles, the ced efficacy against HA-mismatched strains, and the reduced overall efficacy in elderly populations [3,4]. It can be clearly deduced that the heterosubtypic protection against all influenza A viruses can resolve all the above stated setbacks of current influenza A vaccines and prevent the likelihood of epidemics or pandemics from occurring. Immunity to conserve components of a virus may also provide a broad protection against most influenza A strains and subtypes [5,6].

The efficacy of influenza A vaccines designed to induce subtype cross-reactive T-cells to highly conserved internal influenza antigens, such as NP, are exemplified in several animal models using different species [7-10]. DNA vaccination to conserve influenza NP, also known as NP and M, has protected against both matched and mismatched viruses in investigations carried out in both animal and human subjects [9,11-18].

DNA vaccines contain many advantages over conventional vaccines. Live vaccines have proven more advantageous because of their technically simple design, fast formulation period and production time and stability and ease of transport. DNA vaccines also induce strong antigen specific humoral and cellular immune responses against many pathogens such as the influenza virus and tumor antigens [19,20].

The specific humoral and cellular immune responses can be achieved without the use of toxic chemical adjuvants [21]. The DNA vaccine platform presents a promising and effective routs of inducing broad based immunity against influenza. However, frustrations over the efficacy of create platforms need to enhance the potency of DNA vaccines via various optimization methods. There are several feasible approaches available to improve expression and immunogenicity of DNA vaccines in human and animal targets by improving delivery methods, vaccine regimen, and the use of genetic adjuvants. One of the critical roles in this platform is using gene delivery methodology [19,22,23]. Physical delivery such as in vivo electroporation (IEP) significantly increases the transfection efficiency and evokes promising humoral and cellular immune responses in comparison with direct injection of naked DNA [22].

To improve the efficacy of this approach, we explored the ability of a genetic adjuvant fused to NP+M1 vector to enhance the potency of vaccination to NP, an antigen with >90% protein sequence homology among influenza A isolates.

Also, for the first time during this research we fused CRT as a potent immune modulatory molecule that targets the Toll-receptor 4 ligand. The CRT molecule in this experiment displays the promotion of immunogenic APCs function, induces strong CTL responses and prevents induction of tolerance [24].

CRT is a member of the family of HSPs (molecular chaperons) which are conserved and widely distributed in microorganisms and mammalian cells. The main feature of this group of proteins is their profound effect on the immune system. CRT is an endoplasmic reticulum (ER) resident protein that contains a large number of cellular functions is abundantly found in numerous biological systems [25]. CRT is shown to be involved in class I antigen processing and presentation that plays an important role in the formulation of peptide-receptive MHC class I complex [26] and subsequently in efficient peptide loading [24].

CRT demonstrates the ability to bind to peptides transported into the ER by Transporter associated with antigen processing (TAP)[27]. Many studies now indicate that the binding of CRT to peptides may contribute to the production of specific immunity. Immunogenicity is attributed to peptides associated with the extracted CRT and not to the CRT molecule itself. There is also evidence that CRT molecules are capable of being complexed in vitro to unglycosylated peptides and used to elicit a peptide-specific CD8 (+) T-cell response by exogenous administration [28]. Furthermore, when CRT and other members of the HSP family are associated with peptides, they provide a necessary and sufficient source of antigens for cross-priming of CD8 (+) T-cells [29].

In the present study, we constructed a plasmid vector encoding NP-M1-CRT and compared it with NP-M1 sequences with respect to immunogenicity and protective efficacy in a murine model of human influenza A vaccination and virulent challenge. This is the first time that such a study has ever been conducted given its unique approach. In this study, we immunized three groups with just DNA constructs, but for the second group, we primed mice with DNA constructs and boosted them with Live-attenuated influenza virus as a candidate for universal influenza vaccine. The expression of the recombinant vectors was confirmed in the 293T cell line. The potency of this construct to provoke humoral and cellular immune responses and its protective immunity against the lethal challenge of influenza virus infection was then compared with commercially available influenza vaccines and finally evaluated in a murine model system.

## 2. Material and Methods

### 2.1. Ethics statement

All animal care and experiments were conducted in accordance with Institutional Ethic Committee of the Institute of Microbiology, Chinese Academy of Sciences (permit number CASPMI009, CASPMI011). All efforts were made to minimize suffering.

### 2.2. Vaccine Design and Manufacture

The influenza virus utilized in this study was A/PR/8/34 (PR8) supplied by Professor Gao Bin laboratory. Virus stocks were propagated in the allantoic cavities of embryonated hens’ eggs at 37 □C for 24 h.

Cloning of viral structural genes were carried out for NP, M1 and CRT using forward and reverse primers for each construct according to table 1.

**Table 1:**
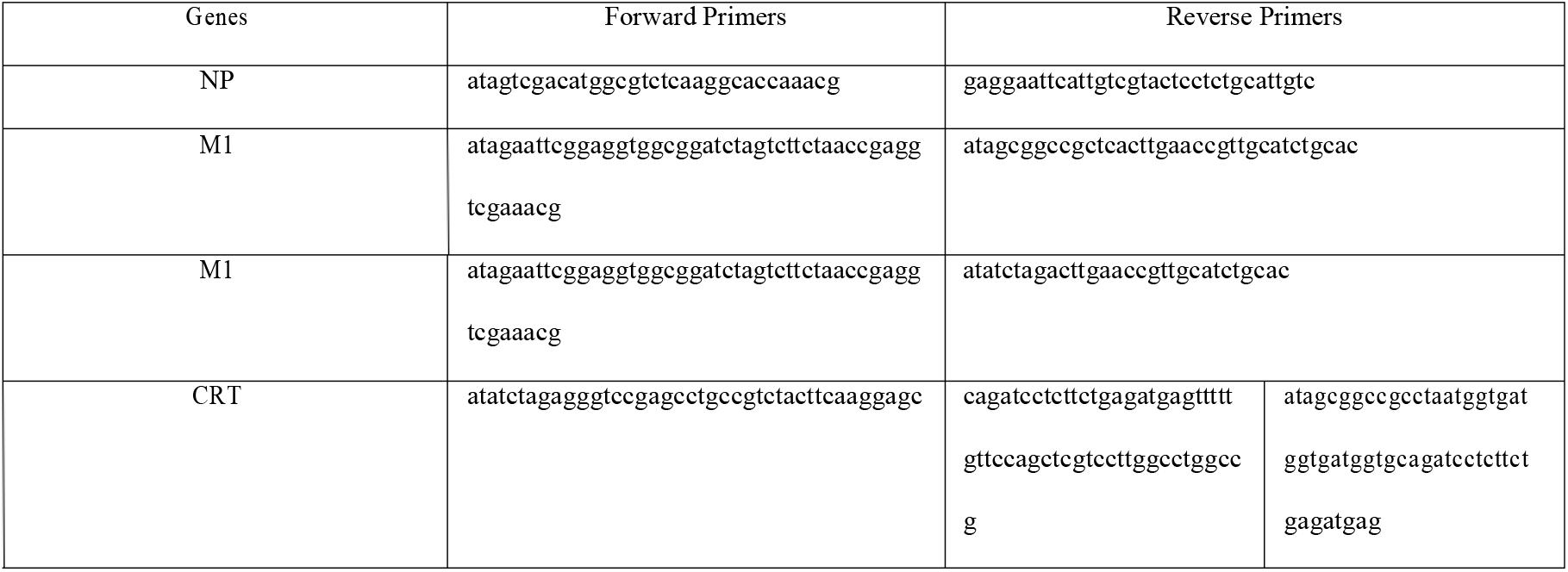
forward and reverse primers for each construct

All sequences were obtained from the National Center for Gene Research, Shanghai Institutes for Biological Sciences.

The vaccine antigens consisted of the complete NP and M1 constructs from the influenza virus PR8 joined by a 7-amino acid linker sequence alone or with CRT as a genetic adjuvant and were expressed in E. coli (BL21). All plasmids were purified according to high purification standards.

The pEF vector used in this study was gifted by Professor Bin Gao (The Center of Molecular Immunology, CAS Key Laboratory of Pathogenic Microbiology and Immunology, Institute of Microbiology, Chinese Academy of Science).

Generation of Live-attenuated influenza virus was performed as described previously in this report [30].

### 2.3. Transient gene expression in mammalian cells

The human embryonic kidney 293T cells (HEK293T) are maintained in Dulbecco’s Modified Eagle’s medium (DMEM) containing 10% fetal bovine serum (FBS). Briefly, one day before transfection, HEK293T from confluent culture were trypsinized and transferred into 10 cm2 plate and cultured for 24 hours (Nunc, Denmark) at 1 × 106 cells per plate. The cells were then incubated at 37 □C in a humidified incubator containing 5% CO2 to achieve approximately 70% confluence on the day of transfection. Calcium phosphate transfection method was used. The cell culture was replaced by Opiti-medium before transfection. Each plate of cells was then transfected with purified expressing plasmids. Six hours after the addition of transfection mixture, 293T cell line was cultured in DMEM medium containing 10% FBS. The cell supernatant was then harvested 48 hours after transfection.

### 2.4. SDS–PAGE and Western-blotting

Forty-eight hours after transfection, the extracts of cell lysates and supernatant were electrophoresed in SDS–PAGE according to Laemmli methodology (Laemmli, 1970). The gel was then stained with Coomassie brilliant blue R-250 and a broad-range marker (Fermentas, China) was used for the estimation of protein size. For western-blotting, separated proteins were transferred onto a polyvinylidene difluoride (PVDF) membrane, blocked by 2% Marvel milk, and then membrane incubated with mouse anti-influenza A virus NP protein monoclonal Ab (Sino biological Company, China; 1: 10,000, 1 h) or mouse anti-influenza polyclonal anti-body (1: 10,000, 1 h). The membrane was then incubated by secondary goat anti-mouse antibody (HRP conjugated, Fermentase; 1: 10,000, 1 h). After three washes, protein bands were detected directly by their exposure to chemiluminescence fluid and radiograph film.

### 2.5. Immunization protocol

Female C57BL/6 mice were purchased from the Division of Beijing HFK Bioscience Ltd. The animals were maintained in a temperature-controlled clean room environment and received food and water ad libitum. Mice were kept for one week in the animal clean room for adaptation and then immunized at 7 weeks of age.

All mice were randomly divided into eight equal groups of 10 at 6-weeks-old (Table 2).

**Table 2:** Experimental design, vaccination status and number of mice in control and treatment groups.

| Group | Vaccination |  |  | Challenge Virus | Total number of mice |
| --- | --- | --- | --- | --- | --- |
| 1 | Vector (Negative control) | Vector (Negative control) | - | PR/8(H1N1) | 10 |
| 2 | NPM1 | NPM1 | - | PR/8(H1N1) | 10 |
| 3 | NPM1CRT | NPM1CRT | - | PR/8(H1N1) | 10 |
| 4 | Split Vaccine | Split Vaccine | - | PR/8(H1N1) | 10 |
| 5 | Vector | Vector | La influenza | PR/8(H1N1) | 10 |
| 6 | NPM1 | NPM1 | La influenza | PR/8(H1N1) | 10 |
| 7 | NPM1CRT | NPM1CRT | La influenza | PR/8(H1N1) | 10 |
| 8 | Split Vaccine | Split Vaccine | La influenza | PR/8(H1N1) | 10 |

La influenza: live-attenuated influenza virus

Mice were immunized beginning at 7 weeks of age (Table 2). DNA vaccination used 100 μ g/mouse in low endotoxin Phosphate-buffered saline (PBS) given i.m. in the quadriceps, half to each leg. They were immunized under lightly anesthesia (10 μ l (6 mg/ml) Pentobarbital sodium/g mouse). Approximately thirty seconds after injection, a pair of electrode patches were attached to the immunization area and the injection sites were electroporated once with the following parameters using an electric pulse generator. Eight 60 V unipolar pulses delivered to each injection site with 200 millisecond intervals with each pulse lasting for 20 milliseconds.

Two doses of DNA vaccine were then given over a duration of two weeks intervals. Two weeks after the last dose of DNA vaccination, three mice in DNA groups were anesthetized by pentobarbital sodium with a dose of 60 μg/g and sacrificed by neck dislocation to study immune responses.Other groups were primed with 2 doses of DNA vaccine and boosted with 30 μ l of Live-attenuated influenza virus (1×104 pfu) intranasally (Table 2). Two weeks after the last immunization, three mice were anesthetized by pentobarbital sodium with a dose of 60 μ g/g and sacrificed by neck dislocation to study immune responses. In the negative control group, mice were immunized with pEF vector.

### 2.6. ELISA

Blood samples for ELISA test were taken prior to immunization and two weeks after immunization. ELISA method is performed as previously described [31]. A total of 96 well plates (Maxisorb, Nunc, Denmark) were briefly coated with 100 μ l of 10 μg/ml NP synthetic peptide solution in 50mM sodium bicarbonate buffer, pH 9.6 and incubated overnight at 4 °C. The plates were then washed with washing buffer and blocked for one hour with PBS-BSA 2% at room temperature. The plates were then loaded with different dilutions of experimental sera and incubated at for two hours at 37 °C. After washing 100 μ l of 1:10000 rabbit anti mouse IgG-HRP conjugates (Sigma-Aldrich), the peptide were added to the wells and incubated for two additional hours at 37 °C. The color reaction developed with 3, 3’, 5, 5’-tetramethylbenzidine, TMB, at OD 450 nm. The optical density (O.D) for NP were measured in experimental groups at different dilution were then compared to each other and with negative control group.

### 2.7. Analysis of virus-specific CTL by tetramer-staining

Mouse CD8_+-_FITC, CD3_+-_PE and H-2D_b-_PE tetramer with the NP_366–374-_ epitope ASNENMETM (Epigen Biotech, Beijing) were used.

Single cell splenocyte suspensions were obtained and red blood cells were removed using erythrocyte lysis buffer. The splenocytes were washed three times with PBS, and 1×105 cells were incubated with CD8_+-_FITC, CD3_+-_PE and H-2D_b--_PE tetramer on ice for thirty minutes. After incubation, the cells were washed twice with PBS. The labeled cells were analyzed by flow cytometry with a FACSCalibur machine (BD Biosciences). All FACS data was analyzed using FlowJo software (Tree Star, Ashland, OR, USA).

### 2.8. Ex Vivo IFN-g ELISPOT

Ex vivo interferon-gamma enzyme-linked immunosorbent spot (IFN-c ELISPOT) assays were performed using mouse spleen cells. The number of NP-specific IFN□ secreting cells in mouse was counted using commercial ELISPOT assay kits (MABTECH AB, Sweden). This procedure was conducted according to the instruction manual. Briefly, 2 × 105 splenocytes were incubated in monoclonal antibody-precoated wells with a mixture of NP peptides (10 μg/ml) for 24 h at 37 °C and 5% CO2. The visible spots of IFN□ secreting cells were enumerated by C.T.L Cellular Technology Ltd. Software.

### 2.9. Viral challenge and follow-up

Two weeks after last immunization, mice were challenged intranasally with a lethal dose (1×107 pfu) of A/PR8 in a total volume of 50 μl.

For challenge experiments, five to six mice per group were used. They were monitored daily for two weeks for mortality and morbidity following monitoring body weight and observation of clinical symptoms at two-day intervals.

### 2.10. Statistical analysis

SPSS 20.0 software (SPSS Inc., Chicago, USA) was used for statistical analysis of the results between treatment and control groups. Data was analyzed for significance (P < 0.05) by the one-way Anova analysis of variance when variance between groups was homogeneous and distribution of the data was normal or a nonparametric test (Kruskal–Wallis) when normality test or homogeneity of variance test failed.

## 3. Result

### 3.1. ELISA Results

Total IgG was evaluated with an in-direct ELISA method. Results show, immunization of mice with DNA constructs and prime-boost groups both induced specific IgG antibody responses against NP peptide which showed significant difference compared with control group in recombinant DNA groups (P≤0.05). Total antibody titer was higher in prime-boost groups but there was no significant difference among these groups (p > 0.05). These results are shown in Figure 1.

**Figure 1.**
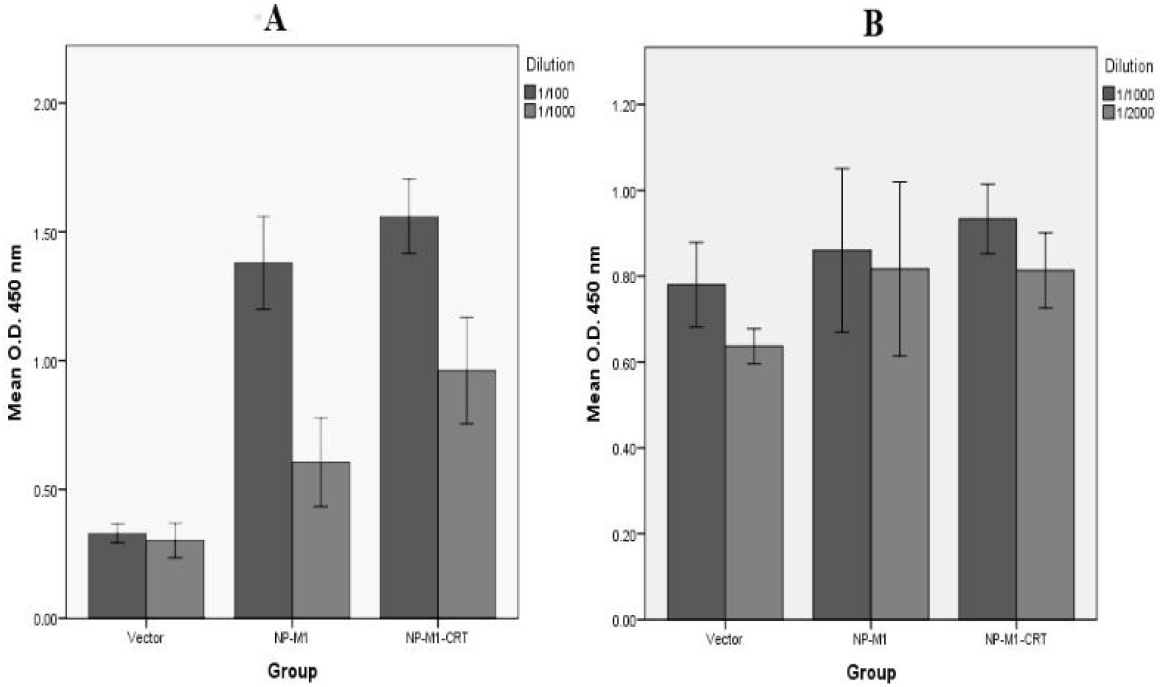
Humoral immune responses against NP peptide in C57/BLC. Anti-sera diluted 1/100 and 1/1000 in DNA construct groups (A). 1/1000 and 1/2000 in prime-boost groups (B). Each bar represents the average antibody responses for each group of immunized mice with standard error (mean ± S.E.M). Mean with different letters between bars are statistically different at the same time point of immunization (P < 0.05, one-way ANOVA).

### 3.2. ELISPOT Results

The IFN-γ secreting lymphocytes were evaluated using ELISPOT assay. As shown in Figure 2, immunization of mice with both recombinant DNA groups and prime-boost groups significantly increased IFN-γ secreting cells as compared to control groups (P≤0.001). High frequency of positive spots appeared only when the spleen cells were stimulated with NP peptide. In contrast, in the negative control group, few spots were observed because the cells could not induce T-cell response.

**Figure 2.**
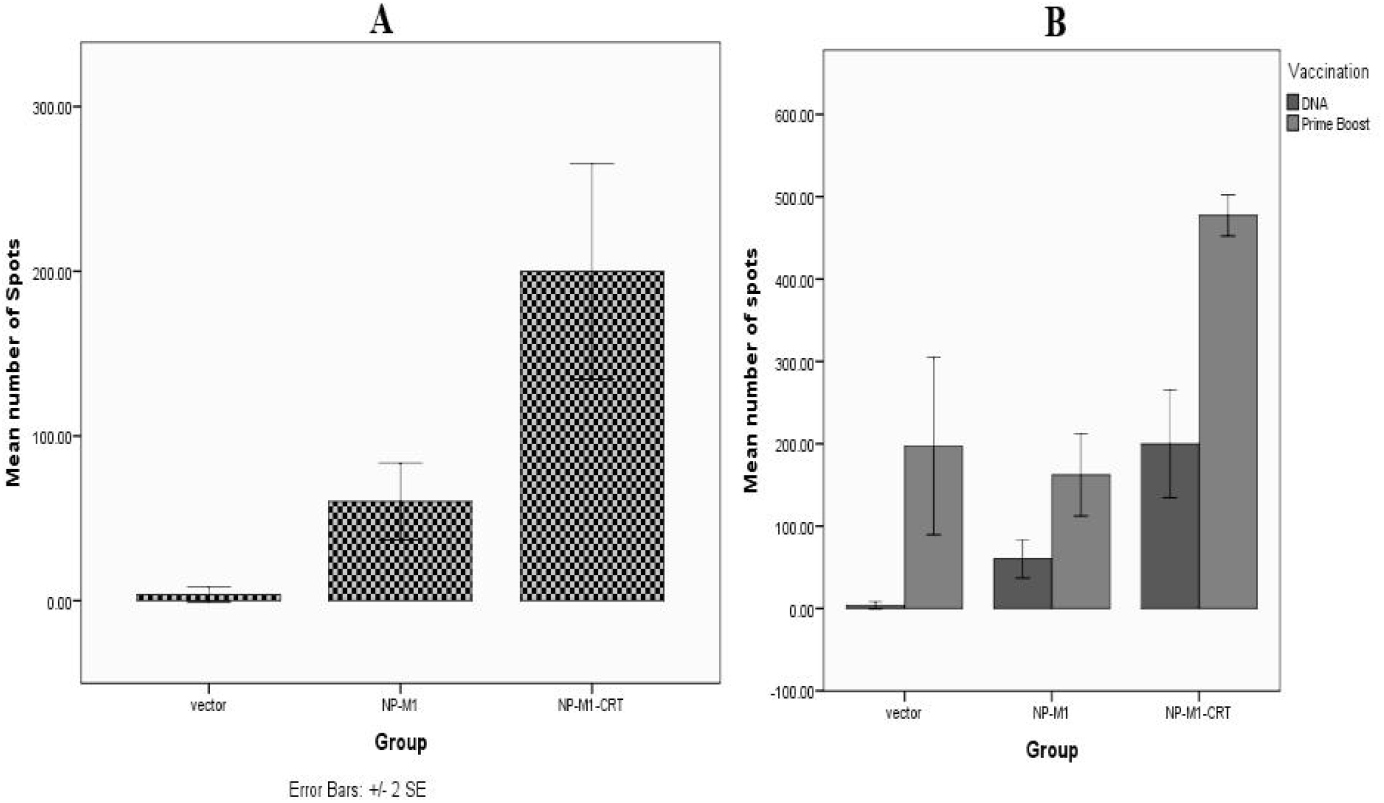
Immunization of mice with recombinant DNA groups (A) and prime-boost groups (B) can activate NP specific T cells. Lymphocytes from the mice immunized with these vaccines are measured by ELISPOT in the presence of 5μg/mL NP peptide. The number of IFN□ secreting lymphocytes is counted. Lymphocytes from un-immunized mice are used as controls. Each bar represents the average for each group of immunized mice with standard error (mean ± S.E.M).

There was no significant difference among NP-M1 and negative control groups in prime-boost immunization. (p > 0.05).

### 3.3. CTL Results

For detection of cytotoxicity of lymphocyte, in vivo CTL assay, mouse CD8_+-_FITC, CD3_+-_PE and H-2D_b--_PE tetramer with the NP_366–374-_ epitope ASNENMETM (Epigen Biotech, Beijing) were used.

Results show that immunization of mice with NP-M1-CRT in both DNA groups and prime-boost groups induced CTL activity as compared with other groups (P≤0.05) although CTL response in prime-boost groups were higher than DNA groups. There was no significant difference among NP-M1 and negative control groups in DNA group immunization. (p > 0.05).

### 3.4. Challenge Results

After challenging the mice with a lethal dose (1×107 pfu) of A/PR8, all groups of the infected mice were evaluated by daily observation to monitor mortality and at two-day intervals for monitoring clinical symptoms and body weight for two weeks. All mice in DNA and prime-boost groups showed weight loss (Figure 4) but in Vector group in DNA immunization on day four to six body weight was less than other groups and clinical signs were clear. In contrast, these clinical signs were delayed for one to two days in mice immunized with NP-M1 and NP-M1-CRT groups in which they also exhibited weight loss and mild clinical signs and then began to recover (Figure 4). The survival rate in the negative control group at 14 days post-challenge was 45%, (Figure 4). The All mice immunized with prime-boost route had enough protection level to show 100% survival rate following challenge with a lethal dose (1×107 pfu) of A/PR8 in contrast with H1N1 split vaccine which had less than 60% survival rate and more weight loss compared with NP-M1 and NP-M1-CRT prime-boost groups.

**Figure 3.**
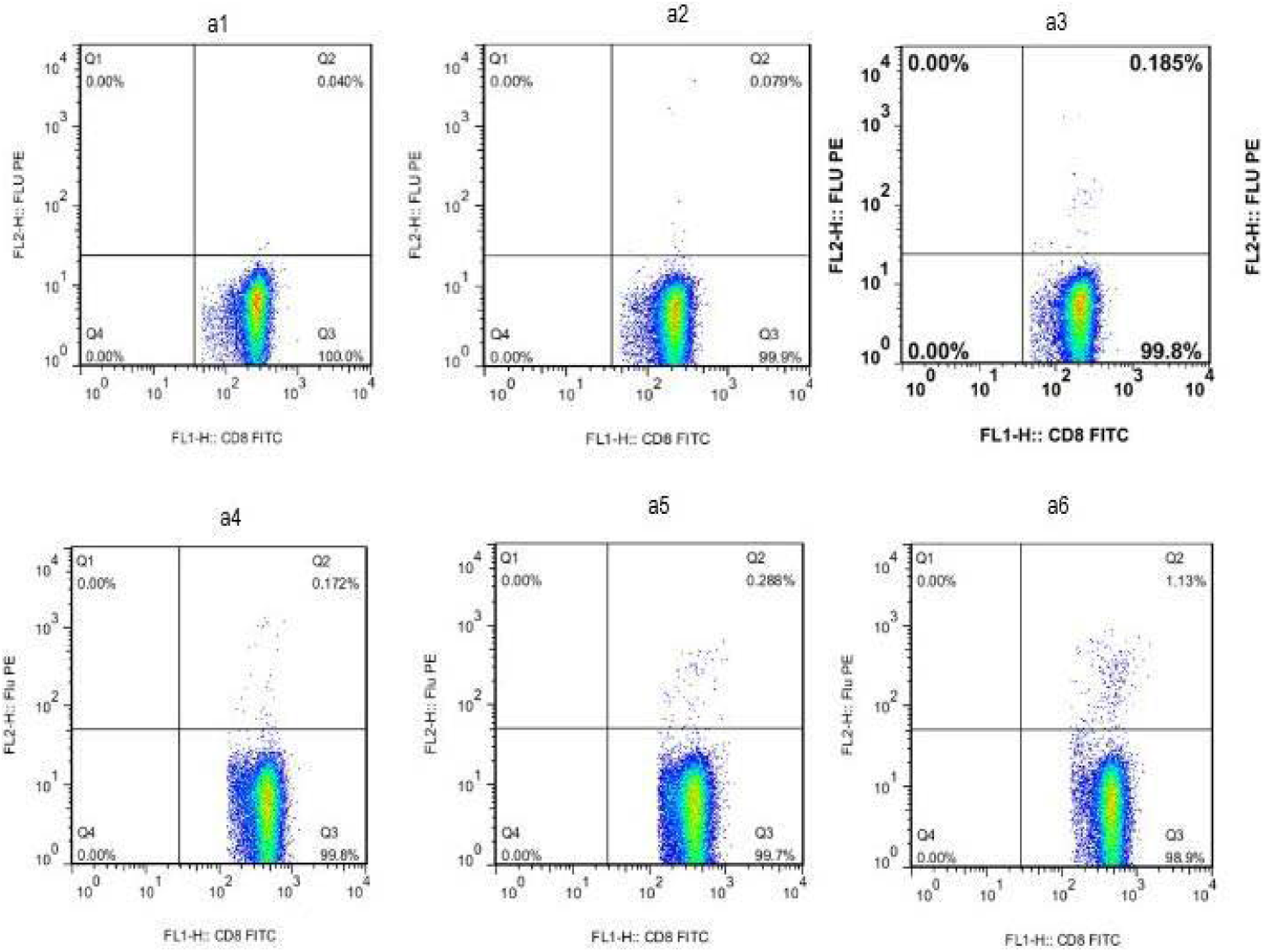
The ratio of CTL responses is determined by flowcytometry. a1: Vector (DNA ONLY), a2: NP-M1 (DNA ONLY) a3: NP-M1-CRT (DNA ONLY), a4: Vector (prime-boost with attenuated influenza virus), a5: NP-M1 (prime-boost with attenuated influenza virus), a6: NP-M1-CRT (prime-boost with attenuated influenza virus).

**Figure 4.**
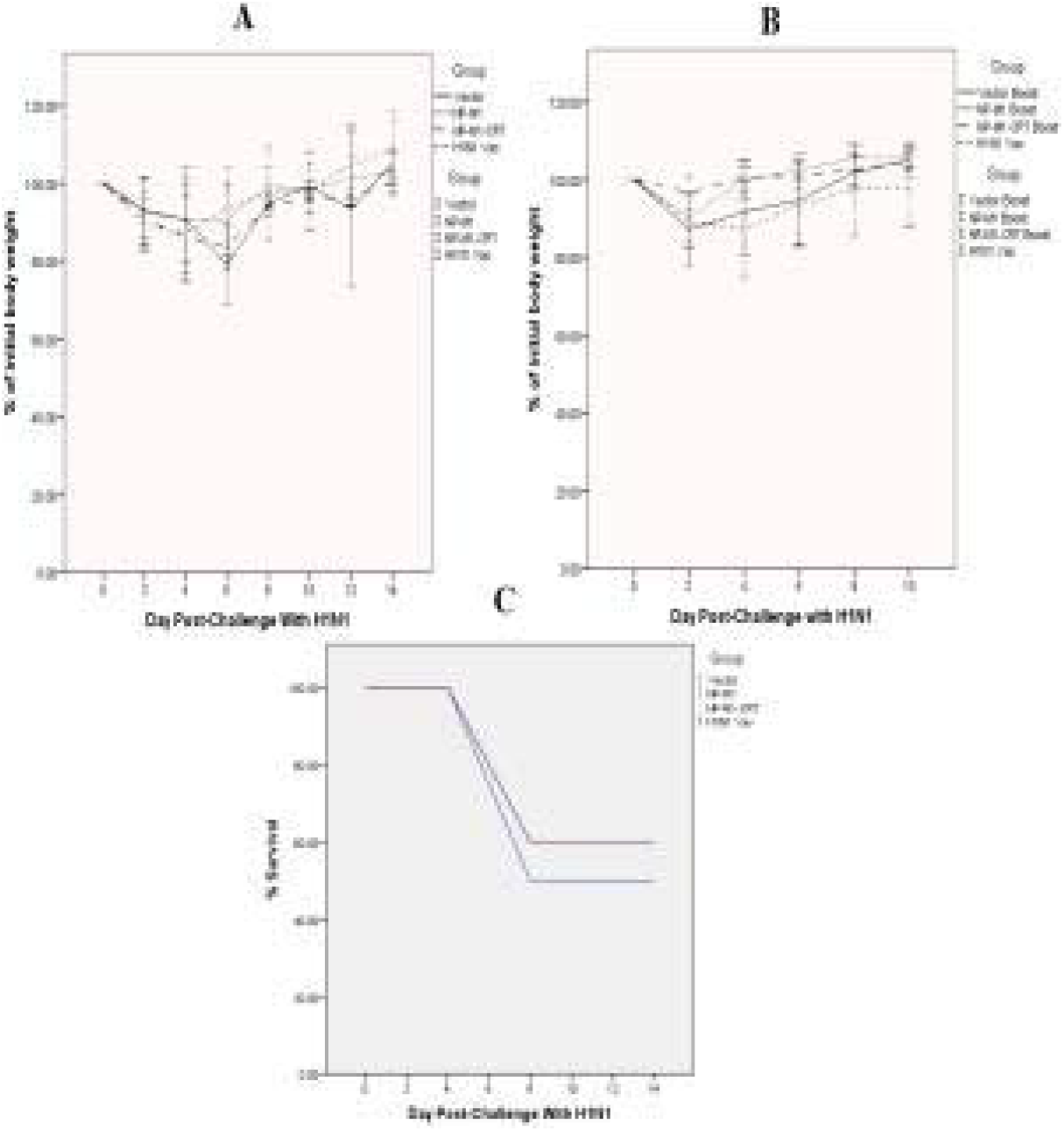
Results of DNA (A) and prime-boost immunization (B) with Vector, NP-M1 and NP-M1-CRT recombinant Constructs alone or in combination with attenuated influenza virus administrated in mice (5 mice/group) and challenges with a lethal dose (1×10^7^ pfu) of A/PR8 influenza A isolates (C). Mice immunized with Vector was challenges as negative control. Survival rate is monitored daily and percent of initial body weight is monitored at two-day intervals for two weeks after viral challenges.

## 4. Discussion

The main drawback of relying on HA as an antigen is the strain specificity of the vaccine [4]. Whereas the conserved NP and M1 proteins of Influenza virus are promising candidate targets for a broad-spectrum, recombinant influenza A vaccine, as they almost completely conserve across all influenza A viruses[1,11]. Some of the earliest research on influenza DNA vaccines show the potency of such a vaccine in different animal models. First, Ulmer et al. (1993) [14] demonstrated that DNA vaccines encoding NP, the conserved internal protein of influenza could protect mice from the heterologous viral challenge. DNA vaccines with different combination of conserved antigens such as M1, M2, NP and HA have shown to be effective in inducing protection to viral challenge in various animal models and human [18,20,32,33].

Luo et al., 2008 [34] demonstrates that when NP fuses to the tissue plasminogen activator (TPA) signal sequence it elicits a strong immune response and potentially can protect mice against H5N1 challenge. Also low immunogenicity against natural NP and M1 may be overcome by fusing them, either genetically or chemically, to an appropriate carrier [11].

Until now, more than six clinical trials as influenza DNA vaccines are being carried out and most of those vaccines are administered parenterally [35, 36]. Another universal influenza DNA vaccine is based on modified vaccinia virus Ankara (MVA), administration of this candidate vaccine (MVA-NP+M1) revealed good safety in humans and induced strong immune responses 6 months after immunization [37,38].

It is important to note that vaccination and gene delivery methods are vital in determining the patterns of cytokines signature and ultimately the mechanisms of protection [39, 40].

CRT carries the potency to motivate polarized cytokines responses toward T helper 1 (Th1), motivate the production of the IgG2a antibody in mice and humans which plays a major role in protection [24,28,29].

To attain an efficient presentation of gene products to immune system, we applied IEP combined with injection of recombinant vector to the quadriceps muscle of the mice. Laddy et al. (2008) displays protection in ferret and mice with IEP against the lethal dose of heterosubtypic influenza A challenge after DNA vaccine encoding the M2e genetically fused to NP [41].

In our study, we have deduced that IEP could be more efficient in inducing higher Th1-type antibody immune responses in contrast with other physical gene delivery methods [40].

To add upon the previously stated point several studies also demonstrate an association between IFN-γ production and the concomitant generation of antigen-specific cytotoxic T-lymphocyte (CTL) responses [42-45].

Virus-specific CTL are important for protection against influenza virus infection [46, 47].

This study also reveals that vaccination with NP-M1-CRT conferred a similar degree of protection from clinical diseases to that induced by the prime-boost vaccination regimen. It also shows that a prime boost vaccination of mice with NP-M1-CRT elicits humoral and cell-mediated immune responses which results in a marked reduction in the severity of clinical disease following challenge with influenza virus. This suggests that NP-specific immune responses can contribute to the protection of mice from influenza infection and disease. Influenza NP is an internal viral structural protein; therefore, NP specific antibodies are not capable of virus neutralization. This might explain the reason why the reduction in duration of viral shedding after prime-boost NP-M1 vaccination was not significantly different from controls (p = 0.228). The inability of NP-specific immune responses to control virus shedding has already previously been reported in other species [48,49] NP typically serves as an important target antigen for cellular immune responses and it can elicit cross-protective immunity to heterologous strains of influenza virus [16,50,51]. Prime-boosted NP-M1-CRT vaccination of mice generates virus-specific lymphoproliferative and IFN-γ responses. Also, a wide variety of influenza virus-specific antibody isotypes were induced following prime-boost NP-M1 vaccination although these responses are typically of lower magnitude than those to HA.

NP peptide is used to coat the ELISA plates for antibody measurement. This further suggests that NP induces quantitatively smaller antibody responses than HA using this vaccination strategy. While NP-M1 DNA alone can protect mice against the low dose virus challenge as previously mentioned in this report); we also show that a DNA prime-live attenuated influenza virus boost regimen is more effective conferring protection against doses of challenge virus that were lethal to mice vaccinated only with DNA. NP-M1-CRT DNA immunization failed to 100% protect against influenza A virus.

Vaccines based on conserved antigens reduce morbidity but do not generally provide sterilizing immunity, which can be viewed as a disadvantage. On the other hand, the limited viral replication they allow can induce additional immune responses, such as anti-H5 antibodies), augmenting subsequent protection.

The lack of boosting after repeated immunizations with the same recombinant virus is probably in part related to the anti-viral antibody and cellular immune response that is primed by the first immunization. In subsequent immunizations, these antibodies may prevent the virus from entering cells and if the boosting dose of the virus succeeds in infecting host cells, the pre-existing response to all the viral antigens will be boosted, rather than only the response to the recombinant antigen. However, priming with DNA and boosting with virus allows the virus to infect cells, express the recombinant antigen and boost the response to the only antigen common to both, resulting in a very high anti-influenza CD8+ T cell response.

In conclusion, the ease of production of these types of vectors and the fact that they would not require annual updating and manufacturing may provide an alternative cost-effective approach to limit the spread of potential pandemic influenza viruses.

Although DNA vaccination combined with in vivo electroporation could significantly improve the immunogenicity of constructed vectors in mice. Also serological evaluations demonstrated the potency of our approach to provoke strong anti-NP specific antibody responses. Moreover, our strategy of immunization in prime-boost groups was able to provide protection against lethal viral challenge using H1N1 subtype. However, further work is needed to evaluate responses to this recombinant vector with different routes of immunization and different delivery methods to make it easier and effective in human.

## Acknowledgments

The project was supported by postdoctoral grants from Chinese Academy of Science. The authors wish to thank Professor Bin Gao (head of CAS Key Laboratory of Pathogenic Microbiology and Immunology, Institute of Microbiology, Chinese Academy of Sciences, Beijing, PR China) and Dr. Mehdi Mahdavi (Virology Dept., Pasteur Institute of Iran) for their helpful contribution to this project.

## Notes

### Competing Interest Statement

The authors have declared no competing interest.

